# Localizing and classifying adaptive targets with trend filtered regression

**DOI:** 10.1101/320523

**Authors:** Mehreen R. Mughal, Michael DeGiorgio

**Affiliations:** Bioinformatics and Genomics at the Huck Institutes of the Life Sciences, Pennsylvania State University, University Park, PA 16802, USA; Departments of Biology and Statistics, Pennsylvania State University, University Park, PA 16802, USA; Institute for CyberScience, Pennsylvania State University, University Park, PA 16802,USA

## Abstract

Identifying genomic locations of natural selection from sequence data is an ongoing challenge in population genetics. Current methods utilizing information combined from several summary statistics typically assume no correlation of summary statistics regardless of the genomic location from which they are calculated. However, due to linkage disequilibrium, summary statistics calculated at nearby genomic positions are highly correlated. We introduce an approach termed *Trendsetter* that accounts for the similarity of statistics calculated from adjacent genomic regions through trend filtering, while reducing the effects of multicollinearity through regularization. Our penalized regression framework has high power to detect sweeps, is capable of classifying sweep regions as either hard or soft, and can be applied to other selection scenarios as well. We find that *Trendsetter* is robust to both extensive missing data and strong background selection, and has comparable power to similar current approaches. Moreover, the model learned by *Trendsetter* can be viewed as a set of curves modeling the spatial distribution of summary statistics in the genome. Application to human genomic data revealed positively-selected regions previously discovered such as LCT in Europeans and EDAR in East Asians. We also identified a number of novel candidates and show that populations with greater relatedness share more sweep signals.

## Introduction

Positive selection is one of the evolutionary processes through which populations adapt to their environments, and identifying positively-selected genomic regions can help us uncover the differences in genes and consequently phenotypes that differentiate populations from one another. Differentiating between diverse types of selective sweeps due to positive selection (Hermisson et al., 2017), such as hard sweeps, which result from a beneficial allele on a single genomic background rising in frequency, and soft sweeps, which occur when a beneficial allele on multiple genomic backgrounds rises in frequency, can also provide us with insights into evolutionary processes. However, identification of adaptive regions is a non-trivial task, as signatures of adaptation are often muddled by demographic events. For instance, both population bottlenecks and selective sweeps can lead to similar decreases in genetic diversity (Wall et al., 2002; Stajich and Hahn, 2004; Jensen et al., 2005). Developments in our understanding of evolutionary mechanisms and their individual importance have led to increasingly complex models (*e.g.*, Nielsen et al., 2005), as well as numerous tests for statistical differentiation between genomic regions undergoing natural selection and neutrality (Vitti et al., 2013).

Several methods have recently been developed that incorporate information from multiple summary statistics to locate positively-selected genomic regions (Lin et al., 2011; Ronen et al., 2013; Pybus et al., 2015; Schrider and Kern, 2016b; Sheehan and Song, 2016; Kern and Schrider, 2018; Sugden et al., 2018). Most existing supervised learning approaches for detecting sweeps use combinations of summary statistics calculated in genomic windows of simulated chromosomes to train classifiers using methods such as support vector machines, random forests, neural networks, and boosting. Differing mechanisms have been employed to handle issues such as missing data and demographic obstruction of selection signatures. For example, the approach taken by Sheehan and Song (2016) attempts to jointly infer demographic and adaptive history. However, this framework requires a tremendous amount of training data, making its application computationally challenging. Schrider and Kern (2016b) use a method of normalizing summary statistics that lessens the impact of demographic events on selection footprints. In both of these approaches, genomic regions missing percentages of data above a certain threshold are not included during analysis, leading to sizable regions labeled as “unclassifiable”.

Current approaches (*e.g.*, Schrider and Kern, 2016b; Sheehan and Song, 2016) attempt to capture the spatial footprint of adaptation by computing summary statistics at adjacent genomic windows. However, such methods do not explicitly account for the autocorrelation expected due to similarity because of physical proximity of these statistics. Regions that have experienced recent selective sweeps due to positive selection exhibit wide stretches of linkage disequilibrium (LD; Kim and Nielsen, 2004; Kim and Stephan, 2002; Sabeti et al., 2002), as recombination has not had sufficient time to erode the signal. Therefore, directly accounting for correlations of summary statistics computed at adjacent genomic regions should be important, and may lead to improvements in the ability to localize adaptive events.

In this article, we introduce a multinomial regression method termed *Trendsetter* that directly models the genomic spatial distribution of summary statistics. We employ trend filtering within a multinomial regression framework to penalize the differences between predictors, constraining them so that they are similar to adjacent values. We explore how penalizing differences in predictors for statistics between one or more adjacent genomic regions transforms the regression model and affects classification. We further compare the performance of *Trendsetter* to leading single-population classification approaches (Lin et al., 2011; Schrider and Kern, 2016b; Kern and Schrider, 2018) developed or modified to differentiate among hard sweeps, soft sweeps, and neutrality. Finally, we apply *Trendsetter* to wholegenome data from worldwide human populations (The 1000 Genomes Project Consortium, 2015), to study the global distribution of sweeps in recent human history.

## Materials and Methods

In this section, we formalize the multinomial regression with trend filtering approach employed by our classifier *Trendsetter*. We discuss choice of summary statistics used as features for the classifier, training and implementation of the classifier, and calibration of class probabilities. We then describe simulation settings and associated parameters to test the performance of *Trendsetter*, as well as its robustness to diverse demographic scenarios, confounding effects of background selection, and missing data. We finalize by discussing the application of *Trendsetter* to empirical data from global human populations.

### Multinomial regression with trend filtering

Trend filtering has enjoyed great attention in a number of fields, including economics (*e.g*., Hodrick and Prescott, 1997), finance (*e.g*., Tsay, 2005), and medicine (*e.g*., Greenland and Longnecker, 1992). The essential idea behind this approach is to fit a non-parameteric curve to time-series or spatially-varying data, in which consecutive data points are highly correlated. Specifically, in the case we consider here, we can imagine that our data points are summary statistics calculated at adjacent single-nucleotide polymorphisms (SNPs), which are correlated due to LD. We would expect that the spatial distribution of statistics calculated at these SNPs should behave like a curve under models of natural selection, in which some statistics are increased or decreased near a site under selection as portrayed in Figure 1.

**Figure 1:**
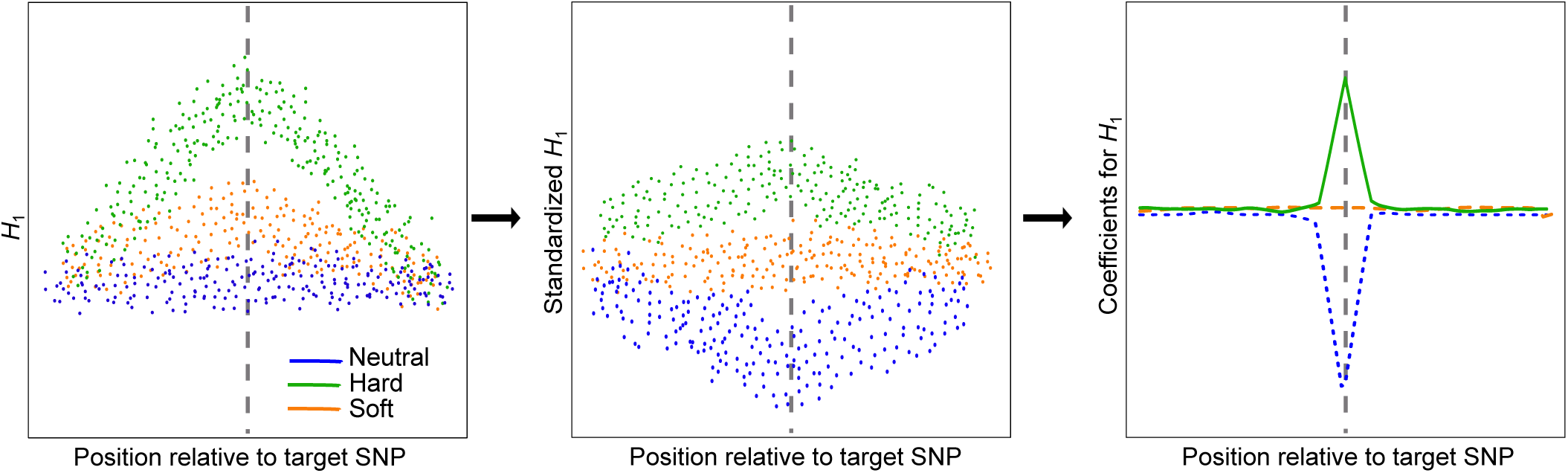
Schematic illustrating steps taken by *Trendsetter* to learn a multinomial regression model. For a given summary statistic (*e.g.*, expected haplotype homozygosity *H*_1_), we compute its value spatially across a genomic region for a set of neutral, hard sweep, and soft sweep simulations used as training data. For *H*_1_, we expect elevated values near the site under selection (target SNP; indicated by a gray vertical dashed line) in sweep simulations, and a greater magnitude of elevation in hard sweep compared to soft sweep settings. This summary statistic is then standardized (mean centered and normalized by the standard deviation) at each position it is computed, so that different summary statistics are comparable. For *H*_1_, this standardization will yield strong negative values for neutral simulations and positive values for hard sweep simulations near a target SNP, and soft sweep simulations will exhibit values intermediate between the neutral and hard sweep scenarios. The model then performs trend filtering on the spatial distribution of each summary statistic (here *H*_1_) for each class (here neutral, soft sweep, and hard sweep), leading to a curve describing the spatial distribution of summary statistics around a target SNP. For *H*_1_, the curve dramatically reduces for the neutral class near the center of the sequence, and is elevated near this position for the hard sweep class.

Here, we plan to perform multinomial regression, accounting for correlations among observations of a particular statistic across neighboring genomic regions through trend filtering. We consider our response to come from *K* classes, and we wish to classify a particular focal SNP as coming from one of the *K* classes. For example, if we have *K* = 3 classes, then we may want to consider responses as neutrality, hard sweep, or soft sweep. To accomplish this task, we will assume that we have observations on *m* summary statistics, with each statistic computed at the focal SNP, and *D* SNP data points upstream and *D* downstream of the focal SNP. These *D* SNPs can either be contiguous or be specified more sparsely across the dataset, which is how we have chosen the set of SNPs in this article. Therefore, for each summary statistic, we will have *p* = 2*D* + 1 observations of the statistic to capture its spatial distribution. We choose to use the spatial distribution of a statistic at SNPs rather than at fixed physical distances (*e.g.*, Chen et al., 2010; Schrider and Kern, 2016b), as it may enhance robustness to missing data when not explicitly accounted for in the training of the classifier.

Suppose we have training data from *n* simulated replicates. Let the true class for simulated replicate *i, i* = 1, 2, *…, n*, be *y*_*i*_. Suppose that the observed value of summary statistic *s* at SNP data point *j* in replicate *i* is denoted by *x*_*i,s,j*_. For observation *i*, denote the probability of observing class *y*_*i*_ given data **x**_*i*_ by ℙ[*y*_*i*_ *|* **x**_*i*_, where **x**_*i*_ is a vector of length *m* × *p* and has transpose

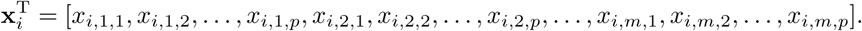

Let *β*_*k,s,j*_ denote the coefficient for class *k, k* = 1, 2, *…, K*, for summary statistic *s, s* = 1, 2, *…, m*, at SNP data point *j, j* = 1, 2, *…, p*. For class *k*, let ***β****k* be a vector of length *m* × *p* that has transpose

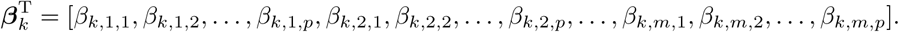

Define the matrix **B** containing *m* × *p* rows and *K* columns by

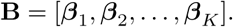

Let *β*_*k,*0_, *k* = 1, 2, *…, K*, denote the intercept for class *k*, and let ***β***_0_ be a vector of length *K* containing these intercept terms with transpose

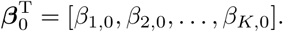

To learn a model that is capable of predicting class *y* from observed data **x**, we need to provide a collection of observed data point tuples 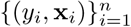 that represent example training inputs **x**_*i*_ and outputs *y*_*i*_ of the model. We then want to learn a model that relates an observed input **x** to an output *y* given model parameters {***β***_0_, **B**}. We therefore wish to compute the conditional probability (Hastie et al., 2009)

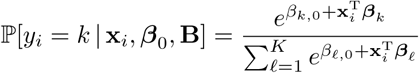

of observing that the output *y*_*i*_ of example *i* was class *k* given the input **x**_*i*_ and model parameters. Given this conditional probability, the log likelihood of the model parameters *{****β***_0_, **B***}* given the collection of observed data point tuples 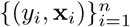 is

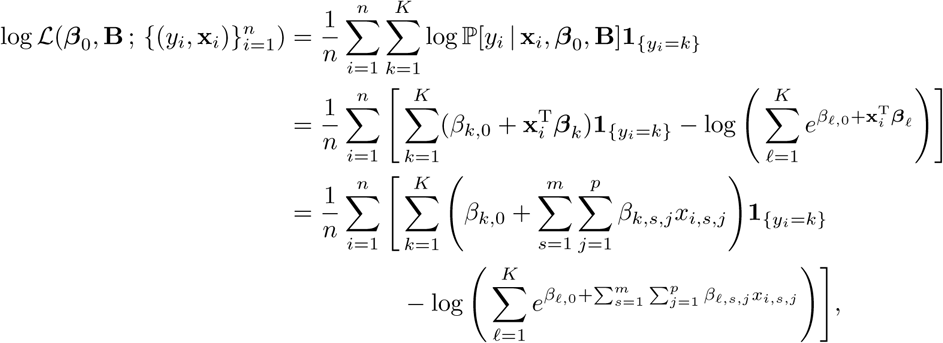

where **1**_{yi=k}_, *k* = 1, 2, *…, K*, is an indicator random variable that takes the value 1 if *y*_*i*_ = *k* and 0 otherwise.

We seek to find the set of coefficients *{****β***_0_, **B***}* that maximize the log likelihood function with a penalty term that we denote PEN_*γ,d*_(**B**), which places a penalty on the coefficients **B**. Denoting the pair of tuning parameters λ_1_ ≥ 0 and λ_2_ ≥ 0, we therefore obtain parameters that maximize a penalized log likelihood function (Hastie et al., 2009) as

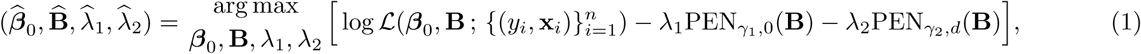

where

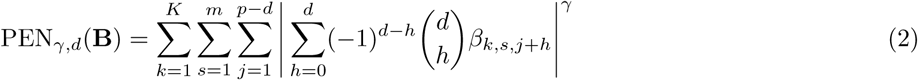

for *γ* ≥ 1 and *d* a non-negative integer. When *d* = 0, 1, or 2, the penalty respectively reduces to

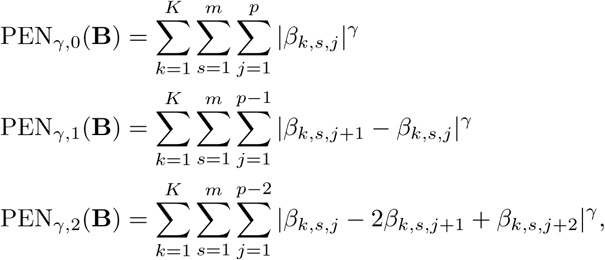

which represent summations across classes and summary statistics for finite difference analogues to the zeroth, first, and second derivatives of functions defined by summary statistic *s* from class *k*. That is, the component of the penalty *β*_*k,s,j*+1_ *-β*_*k,s,j*_ in the second equation (*d* = 1) represents an approximation to the first derivative of the function defined by statistic *s* at SNP data point *j* for class *k*, whereas the component of the penalty *β*_*k,s,j*_ -2*β*_*k,s,j*+1_ +*β*_*k,s,j*+2_ in the third equation (*d* = 2) represents an approximation to the second derivative of the function defined by statistic*s* at SNP data point *j* +1 for class *k*. In general, 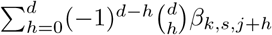 represents a finite difference approximation to the *d*th derivative of the function defined by statistic *s* for class *k* at SNP data point *j*. Letting *γ* = 1 gives the *l*_1_ penalty commonly employed in lasso, and setting *γ* = 2 gives the *l*_2_ penalty commonly used in ridge regression frameworks (Hastie et al., 2009).

In this article, we consider the situation in which *γ*_1_ = *γ*_2_ = 1, permitting simultaneous regularization and feature selection. The first penalty term PEN_1,0_(**B**) associated with tuning parameter λ_1_ is identical to the one used by lasso (Hastie et al., 2009). This penalty ensures that the values of regression coefficients for summary statistics that are highly correlated with other selected (important) summary statistics will be reduced to zero, thus reducing the effects of multicollinearity. In contrast, the second penalty term deals with the autocorrelation of summary statistics, or how each summary statistic is correlated across physical space. For the second penalty term PEN_1,*d*_(**B**) associated with tuning parameter λ_2_, we consider values of *d* = 1 and *d* = 2. The scenario with *d* = 1 approximates a function with a step or piecewise-constant function and is termed constant trend filtering, whereas *d* = 2 approximates a function with a piecewise-linear function and is termed linear trend filtering (Kim et al., 2009; Hawkins and Maboudou-Tchao, 2013; Tibshirani, 2014; Wang et al., 2016). Using *d* = 2 measures the curvature of the function for statistic *s* at SNP data point *j* +1. The entire penalty PEN_1,2_(**B**) therefore represents the total curvature across all summary statistics, and assesses the ruggedness of the set of curves. By penalizing in this manner, we are imposing a smoothness on the spatial distribution of the summary statistics. The combination of this trend penalty with that of the lasso penalty PEN_1,0_(**B**) has a similar effect to a group lasso (Ming and Yi, 2006) penalty, in which the inclusion or exclusion of all values of a summary statistic is decided rather than the inclusion or exclusion of each feature separately. Other trend penalties focusing on lower- and higher-order derivatives have been considered in the literature (Tibshirani, 2014; Wang et al., 2016).

### Choosing summary statistics

The choices of summary statistics are critical when designing a regression approach for isolating signals of natural selection. First, summary statistics that interrogate different aspects of genetic variation are important. For example, statistics such as the mean pairwise sequence difference 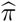 (Tajima, 1983) can be used to evaluate skews in the site frequency spectrum. Linkage disequilibrium statistics, such as the squared correlation coefficient *r*^2^ (Hill and Robertson, 1968) between a pair of SNPs can be used to evaluate speed of decay of SNP correlation with distance from a focal SNP. Furthermore, summaries of haplotypic variation, such as the number of distinct haplotypes *N*_haps_ and expected haplotype homozygosity *H*_1_ (Garud et al., 2015) can be used to evaluate skews in the distribution of haplotypes as a function of distance from a focal SNP. Second, summary statistics that should be relatively robust to the confounding effects of background selection, such as haplotype-based statistics (Enard et al., 2014), should be considered, as background selection has been demonstrated to be a ubiquitous force in a number of diverse lineages (*e.g.*, McVicker et al., 2009; Comeron, 2014). In this article, we focus on a set of *m* = 6 summary statistics, including mean pairwise sequence difference 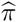(Tajima, 1983), the squared correlation coefficient *r*^2^ (Hill and Robertson, 1968) of a SNP and the focal SNP, the number of distinct haplotypes *N*_haps_, and the *H*_1_, *H*_12_, and *H*_2_*/H*_1_ statistics of Garud et al. (2015). The latter three statistics were chosen as they have been demonstrated to exhibit high power to detect both hard and soft sweeps, as well as the ability to collectively distinguish between hard and soft sweeps.

It is important to note that it is possible to extend our approach to unphased genotypes, by using unphased-genotype analogues of the haplotype-based statistics. That is, following Harris et al. (2018), we can substitute the number of distinct haplotypes *N*_haps_ with the number of distinct multilocus genotypes *N*_geno_, replace *H*_1_, *H*_12_, and *H*_2_*/H*_1_ respectively with *G*_1_, *G*_123_, and *G*_2_*/G*_1_ (Harris et al., 2018), and use *HR*^2^ (Sabatti and Risch, 2002) as a surrogate for *r*^2^. Because the multilocus genotype analogues *G*_1_, *G*_123_, and *G*_2_*/G*_1_ have been demonstrated to retain similar detection and classification abilities as *H*_1_, *H*_12_, and *H*_2_*/H*_1_ (Harris et al., 2018), they should be suitable substitutions. Making these summary statistic substitutions permits application of *Trendsetter* to data from organisms that cannot be phased, as well as for studies in which it is important to avoid phasing errors (see *Discussion*).

### Training the classifier

We computed the value of a summary statistic at each of the 2*D* +1 SNP data points, as described in the *Multinomial regression with trend filtering* subsection. Specifically, for each SNP data point, we considered five SNPs directly upstream and five SNPs directly downstream of the SNP data point, making a window of 11 total SNPs (Figure S1). Each summary statistic was calculated using the data across these 11 SNPs. Specifically, *N*_haps_, *H*_1_, *H*_12_, and *H*_2_*/H*_1_ for a given SNP data point were based on the haplotypic variation defined by the 11 SNP window surrounding (and including) the SNP data point. The mean pairwise sequence difference 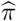 for a given SNP data point was computed as the mean across all 11 SNPs in the window surrounding (and including) the SNP data point. The squared correlation coefficient *r*^2^ for a given SNP data point was computed as the mean *r*^2^ for all 11 SNPs in the window with the focal SNP within the set of 2*D* + 1 SNP data points. Computing *r*^2^ in such a way permitted the method to evaluate the speed at which LD decays from a focal SNP data point (putative site under selection). In this article, we use *D* = 100, so that each summary statistic is computed across 201 data points. Adjacent SNPs will be highly correlated, and we have a trade-off between the number of data points to learn the function for the summary statistic through trend filtering and the running time due to increased numbers of features. To accomplish this, we chose to compute SNP data points every five SNPs, so that we still capture the genomic signal across a wide spatial distribution, while also having adjacent data points that are highly correlated. Such an approach permits us to examine the spatial variation of a summary statistic spanning a total of 10(*D* + 1) SNPs, while only using 2*D* + 1 data points. As a consequence of how we compute summary statistics at each data point, values at neighboring data points will be based on data from a partially overlapping set of SNPs, and will therefore be correlated by construction. For a sample of size 100 haplotypes, in a population with diploid effective size *N* = 10^4^ (Takahata, 1993) and per-site per-generation mutation rateμ = 1.25 × 10 ^-8^ (Scally and Durbin, 2012), setting the 10(*D* + 1) = 1010 SNPs (segregating sites) equal to its neutral expectation (Ewens, 1974) gives the expected length of a neutrally-evolving region with this many SNPs to be 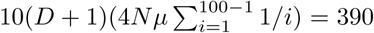, 159 nucleotides, or approximately 390 kb. In regions that have undergone recent strong selective sweeps, much of the genetic variation would have been lost, and so the genomic region with the same number of SNPs would be considerably wider.

To train a classifier under a given demographic model, we used the coalescent simulator discoal (Schrider and Kern, 2016a) to generate 10^3^ neutral, 10^3^ hard sweep, and 10^3^ soft sweep scenarios to use as training data, and computed the 2*D* + 1 values for each of the six summary statistics using the set of SNPs closest to the center of the simulated region. All simulations assumed a uniform per-site per-generation mutation rate of μ = 1.25 × 10^-8^ (Scally and Durbin, 2012) and a uniform per-site per-generation recombination rate of *r* = 10^-8^ (Payseur and Nachman, 2000) across sequences of length *L* = 1.1 Mb. For all selection simulations, beneficial mutations were introduced at the center of the simulated region with per-generation selection coefficient *s* drawn uniformly at random on a log scale over the interval [0.005, 0.5]. Moreover for soft sweeps on standing variation, the starting frequency of the beneficial allele was drawn uniformly at random over the interval [0.01, 0.10]. For all selective sweep simulations, the time at which the adaptive allele reached fixation was drawn uniformly at random between zero and 1,200 generations in the past. Note that because discoal conditions on the time at which a sweep completes, given a specified selection coefficient, the time at which the sweep initiated is already some function of these parameters. Therefore, sweeps associated with small selection coefficients tend to initiate farther in the past than those with larger selection coefficients. Moreover, because discoal allows the specification of both sweep strength and sweep completion time, selected alleles with very small selection coefficients will still not be lost, but will instead have been introduced distantly in the past. As a consequence, sweeps from such weakly-beneficial alleles would likely not be detectable by *Trendsetter*.

As is common for regularized regression models (Hastie et al., 2009; Simon and Tibshirani, 2012), values for summary statistic *s* at SNP data point *j* were standardized so that they had mean zero and standard deviation one across the set of 3 × 10^3^ simulated training replicates. We then used Equation 1 to estimate the coefficients from these data points using 10-fold cross validation (Hastie et al., 2009) with balanced training samples from each class. We subsequently applied *Trendsetter* to simulated and empirical data to classify focal SNPs, where we standardized each summary statistic in the test and empirical datasets using the standardization parameters we applied for the respective training sets.

## Implementation

The optimization problem in Equation 1 is convex, but is non-trivial as it contains two different components—one that is smooth (*i.e.*, the log likelihood function) and the other that is non-smooth (*i.e.*, the penalty function). Liu et al. (2010) developed an efficient algorithm for solving this problem, and we adapted this framework for our purposes. Specifically, we augmented the approach of Liu et al. (2010) to add linear trend filtering, which requires solving a pentadiagonal rather than tridiagonal system of linear equations as was used by the original constant trend filtering implementation. More generally, for a given value of the derivative *d*, this linear system amounts to inverting a symmetric banded Toeplitz matrix with bandwidth *d*. To ensure that the optimization is computationally feasible in reasonable time, we employed the PTRANS-1 algorithm (Liu et al., 2010) for solving general pentadiagonal linear systems, which requires only *O*(*n*) operations for a matrix of size *n* × *n* (where *n* = *p -* 2 = 2*D -* 1 in our scenario for linear trend filtering), and therefore has complexity *O*(*D*) for *D* SNP data points flanking either side of the focal SNP.

## Calibrating class probabilities

Our model, similar to others (*e.g.*, Lin et al., 2011; Schrider and Kern, 2016b; Sugden et al., 2018), not only assigns class labels, but also provides a probability for each of the *K* classes. A properly-calibrated classifier should be one in which the probability of observing a given class is the actual fraction of times that the classifier chooses this class. This calibration ensures that the assigned probability for each class can be interpreted as the empirical proportion of simulations at each threshold.

To examine whether *Trendsetter* yielded properly-calibrated probabilities, we plot a set of reliability curves for each trained classifier. To calibrate classifier probabilities, it is possible to employ Platt scaling (Platt, 1999) applied to the output probabilities of a classifier. Specifically, an extra set of training data must be set aside to train a multinomial logistic model using the probabilities output from *Trendsetter* as the independent variables, and the true class as the dependent variable (Naeini, 2017). To calibrate our classifiers, we use 1000 examples from each class, for a total training set of 3000. Because we expect the majority of polymorphisms in some species (*e.g.*, humans) to be classified as neutral, it may in some cases be useful to calibrate a classifier with this assumption in mind, as increasing the number of neutral examples may be necessary to achieve proper calibration (Sugden et al., 2018).

### Simulations to examine *Trendsetter* performance

We examined a number of simulation settings to better understand the ability of *Trendsetter* to detect and classify sweeps, as well as its robustness to common confounding factors. Specifically, we considered differences in demographic history inspired by population size fluctuations inferred from human genomic data (Terhorst et al., 2017), the influence of soft shoulders (Schrider et al., 2015), background selection due to long-term purifying selection (McVicker et al., 2009; Comeron, 2014), extensive missing data due to regions of poor alignability or mappability, sample size, and selection strengths.

### Demographic history

We considered a constant-size demographic history with effective size of *N* = 10^4^ diploid individuals (see *Training the classifier*), as well as models incorporating population size change that are inspired by parameters inferred from human history (Terhorst et al., 2017)—with models that incorporate recent population expansions that occurred in populations of sub-Saharan African ancestry (*e.g.*, LWK and YRI), and models with strong recent population bottlenecks that occurred in populations of non-African ancestry (*e.g.*, GIH, TSI, CEU, CHB, and JPT). We used these piecewise-constant demographic histories inferred by Terhorst et al. (2017) to train our models. We chose to use the models of Terhorst et al. (2017) instead of those from Tennessen et al. (2012), because in addition to allele frequency information used by Tennessen et al. (2012), Terhorst et al. (2017) also incorporated patterns of LD to infer demographic histories, thereby potentially making their inferred models more accurate (Beichman et al., 2017)— though linked-selection may bias inferences of demographic history from whole-genome methods that incorporate LD, and so masking such regions, as can be done within the Terhorst et al. (2017) framework, may be important. We used 200 time points and corresponding effective population sizes throughout human history for each of our seven populations of interest, which included African (YRI and LWK), European (CEU and TSI), South Asian (GIH), and East Asian (CHB and JPT) groups (see *Application to empirical data*). We utilized these data points as 200 intervals describing the growths and declines of these populations as inputs to discoal along with a range of selection strengths s ∈ [0.005, 0.5] for hard and soft sweeps. The per-site per-generation mutation and recombination rates used for simulating all Terhorst et al. (2017) demographic histories are *μ* = 1.25 × 10^-8^and *r* = 10^-8^, respectively. In simulations with hard and soft sweeps, we ensure the beneficial allele fixes between 1200 generations ago and the present, with fixation time drawn uniformly at random over this time period.

### Linked-sweep classes

Previous work has shown that when classifying genomic regions with window-based methods, it may be possible to mis-classify genomic regions near a hard sweep as soft sweeps via a phenomenon termed “soft shoulders” (Schrider et al., 2015; Schrider and Kern, 2016b). To test whether *Trendsetter* is affected by soft shoulders, we simulated linked-sweep regions by moving the location of a beneficial mutation away from the center by steps of 100 kb, in both the upstream and downstream directions. We do this for both hard sweeps and soft sweeps. To form the training set for the linked sweep classes, we combine 100 simulations from each of the 10 sets of sweep simulations with selected sites distant from the test site.

### Background selection

To evaluate the robustness of *Trendsetter* to regions evolving under background selection, we first followed the protocol described in Schrider and Kern (2017). We employed the forward-time simulator SLiM 2 (Haller and Messer, 2017) to generate 10^3^simulated replicates for sequences of length 1.1 Mb, where the number, lengths, spatial distribution, and distribution of fitness effects of functional elements across the simulated region matched random regions from the human genome. Specifically, we sampled a 1.1 Mb region of the human genome uniformly at random, and determined the sites within that region that are either included in the phastCons database (Siepel et al., 2005) or found within an exon in the GENCODE database (Harrow et al., 2012). In simulations, sites falling within these regions were determined to be undergoing purifying selection, with 25% of mutations occurring in these sites being neutral and 75% having a selection coefficient drawn from a gamma distribution with mean -0.0294 as described by Boyko et al. (2008).

Along with this empirically-based background selection scenario, we wanted to investigate a potentially more extreme setting, with a single genic element located at the center of a large genomic region, in which strongly deleterious alleles arise continually within this genic element. We also used SLiM 2 to generate 10^3^simulated replicates for sequences of length 1.1 Mb, where a central 11 kb “gene” evolved under purifying selection. Specifically, this central genic region was composed of 5’ and 3’ untranslated regions (UTRs) that flanked a set of 10 exons, which were separated by introns as in Cheng et al. (2017). We set the lengths of these UTRs, exons, and introns to be based on their means in the human genome (Mignone et al., 2002; Sakharkar et al., 2004), such that the lengths of each intron, exon, 5’ UTR, and 3’ UTR were 1000, 100, 200, and 800 nucleotides, respectively. We simulated differences in proportions of deleterious mutations arising in each of these genic elements, by simulating 75%, 50%, and 10% of mutations arising in exons, UTRs, and introns as deleterious, respectively, and deleterious mutations having a strong selective disadvantage of *s* = -0.1 per generation. Finally, as a third background selection scenario, we also considered the exact setting as this second central genic element scenario, but with the recombination rate decreased to 100-fold lower in the 11 kb genic region relative to the surrounding neutral regions. This scenario permitted us to examine whether *Trendsetter* was robust to strongly-deleterious mutations arising in regions of elevated linkage disequilibrium. We then tested whether this set of three background selection settings would be falsely classified as a sweep by *Trendsetter* trained using simulations of the constant-size demographic history discussed in section *Training the classifier*.

### Missing data

Due to a number of technical issues, large segments of missing data are scattered throughout the genome (Lander, 2011). Filtering such segments can lead to a large fraction of the genome that cannot be classified (Schrider and Kern, 2016b; Sheehan and Song, 2016), unless it is properly accounted for within the training dataset, as missing data can masquerade as footprints of lost diversity, mimicking patterns expected from selective sweeps. *Trendsetter* computes summary statistics at SNP data points rather than as averages over physical regions, potentially enabling it to be robust to falsely attributing regions with missing data as candidate sweeps. The rationale is that missing data would cause SNP data points to be farther in terms of both physical and genetic distance than if there was no missing data, thereby making SNP data points close to the focal SNP less correlated than expected under selection models. Such an approach should be conservative, likely leading to classifications of sweeps as neutral and not mis-classifying neutral regions as sweeps. To evaluate robustness to missing data, we masked SNPs in the testing dataset, amounting to 30% of the total number of SNPs in each simulation and approximately 30% of the total length of the chromosome. We did this by removing 10 genomic chunks each with size equalling 3% of the total number of simulated SNPs, with a starting position for each missing chunk chosen uniformly at random from the set of SNPs, provided the chunk did not overlap with previously-missing chunks. Removing SNPs in this fashion simulates missing data that would be filtered due to genomic regions with poor alignability or mappability (Mallick et al., 2009).

### Effect of sample sizes on classification rates

The number of individuals sequenced can differ in projects depending on sample availability and funding resources for sequencing. Larger sample sizes are expected to yield better estimates of summary statistics, and therefore more accurate interrogations of genomic diversity. To explore how sample size affects classification accuracy, we tested the ability of *Trendsetter* to correctly classify hard sweeps, soft sweeps, and neutral regions as a function of sample size, choosing sample sizes of 100, 25 and 10 diploid individuals for a set of selection strengths *s* ∈ [0.005, 0.5] ranging from moderate to strong.

#### Selection strengths and classification rates

The strength of selection has an impact on the speed at which a selected allele increases in frequency toward fixation, and thus the amount of time for mutation and recombination to erode the signature. Specifically, the size of the genomic footprint *L*_footprint_ can be approximated by the equation *L*_footprint_ = *s/*(2*r* ln(4*Ns*)), where *s* is the per-generation selection coefficient, *r* is the per-site per-generation recombination rate, and *N* is the diploid effective population size (Gillespie, 2004; Hermisson and Pennings, 2005; Garud et al., 2015). Here, the footprint is positively correlated with the strength of selection, whereas it is negatively correlated with the rate of recombination. To test the effects that different selection strengths have on overall classification rates, we simulated hard and soft sweeps with selection coefficients chosen from two non-overlapping intervals: strong selection with *s* ∈ [0.05, 0.5] and moderate selection with *s* ∈ [0.005, 0.05]. We conducted simulations under a constant-size demographic model using discoal as described in section *Training the classifier*.

#### Application to empirical data

We used phased haplotypes from variant calls of the 1000 Genomes Project (The 1000 Genomes Project Consortium, 2015). Specifically, we analyzed genomes from the sub-Saharan African Yoruban (YRI) population, Gujurati Indian from Houston, Texas, USA (GIH), Han Chinese in Beijing, China (CHB), Japanese in Tokyo, Japan (JPT), Luhya in Webuye, Kenya (LWK), Toscani in Italy (TSI) and Utah Residents with European Ancestry (CEU). We first filtered regions with poor mappability and alignability as in Huber et al. (2016). Specifically, we segmented each chromosome into 100 kb non-overlapping regions, and filtered SNPs in regions with a mean CRG100 score (Derrien et al., 2012) less than 0.9. Because sweeps tend to affect large genomic regions, filtering in this manner will remove large regions with poor average quality, decreasing the likelihood that *Trendsetter* would be misled by genetic variation in unreliable genomic regions. After masking these regions, we computed summary statistics in an identical manner as for the simulated datasets. However, for each chromosome, we classify every fifth SNP beginning from the 505th using information from 100 data points (505 SNPs) upstream and 100 data points (505 SNPs) downstream of the focal SNP. It is important to note that each focal SNP classified by *Trendsetter* should not be viewed as the site or exact location of the beneficial mutation, but rather should be considered as a proxy for the location. If the focal SNP is close enough to the location of the adaptive variant, then we would expect the genetic variation surrounding the focal SNP to look similar to the diversity around the adaptive site. Therefore, using summary statistics that compute a single value at a given SNP (*e.g.*, iHS or XP-EHH; see *Discussion* section) may enhance the ability of *Trendsetter* to only classify polymorphisms that are close to the true adaptive site as a sweep.

When training *Trendsetter* for application to empirical data, our treatment differed from when we evaluated *Trendsetter*’s performance on simulated data in two ways. First, because human recombination rate varies across the genome, we accounted for recombination rate variation by following Schrider and Kern (2017) and drawing the recombination rate for a particular training simulation from an exponential distribution with mean 10^-8^and truncated at three times the mean. Second, because we have introduced another variable into our training simulations (namely recombination rate), we chose to increase the number of independent training simulations by five fold, leading to 5000 neutral, 5000 hard sweep, and 5000 soft sweep replicates.

#### Comparison to other methods

A number of powerful approaches have recently emerged to localize and classify sweeps from genomic data. We compare the classification ability of *Trendsetter* to the binary classifier evolBoosting (Lin et al., 2011), as well as the multi-class approaches of S/HIC (Schrider and Kern, 2016b) and diploS/HIC (Kern and Schrider, 2018). Following Schrider and Kern (2016b), to compare binary to multi-class classifiers, we expanded evolBoosting to greater than two classes by training a classifier to differentiate between sweeps (combined hard and soft) and neutrality, and training another classifier to differentiate between hard and soft sweeps. This procedure is analogous to how Lin et al. (2011) differentiated among sweeps, population bottlenecks, and a constant-size demographic history in the article that introduced evolBoosting. Moreover, to enable direct comparison of S/HIC and diploS/HIC to *Trendsetter*, we employed three-class versions of S/HIC and diploS/HIC approaches, whereas their native states include five classes. We later expand *Trendsetter* to five classes to permit direct comparison with the default states of S/HIC and diploS/HIC. In addition to direct comparison across methods, we also evaluated detection capabilities and robustness to confounding factors when *Trendsetter* operates on the expanded set of summary statistics used by S/HIC. Specifically, S/HIC uses 10 summary statistics: Tajima’s *D* (Tajima, 1983), the maximum value of ω (Kim and Nielsen, 2004), Tajima’s 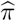(Tajima, 1983), *H*_1_ (Garud et al., 2015), *H*_12_ (Garud et al., 2015), *H*_2_*/H*_1_ (Garud et al., 2015), number of haplotypes *N*_haps_, *Z*_*ns*_ (Kelly, 1997), Fay and Wu’s *H* (Fay and Wu, 2000), and Watterson’s 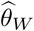 (Watterson, 1975) calculated in each of 11 contiguous windows. Because *Trendsetter* uses many more data points for summary statistics to capture their spatial distribution across the genome, we computed each of the 10 summary statistics in each of 110 contiguous windows, where each window was 1/10th the size of the window used by S/HIC, thereby requiring *Trendsetter* to operate on the same data.

We tested the classification rates for both the constant (*d* = 1) and linear (*d* = 2) trend penalties employed by *Trendsetter*. Moreover, because S/HIC was developed to classify genomic regions as either neutral, a sweep, or linked to a sweep, we also included linked-hard and linked-soft classes to examine whether they enhance the robustness of *Trendsetter* to soft shoulders (Schrider et al., 2015; Schrider and Kern, 2016b). Finally, we compared *Trendsetter* to diploS/HIC (Kern and Schrider, 2018), a recently-developed approach that utilizes deep neural networks and image analysis to learn the spatial distribution of summary statistics nearby a sweep region—similar in concept to accounting for the spatial orientation of summary statistics that gives *Trendsetter* its power. Similarly to testing with S/HIC-specific statistics, we tested *Trendsetter* using the statistics specified by diploS/HIC in 110 contiguous windows. This feature vector includes statistics measuring the variance, skewness, and kurtosis of the distribution of multilocus genotype distances.

### Results

To examine the power and robustness of *Trendsetter*, we evaluate its performance under common settings that would typically be encountered in empirical data. Specifically, we test the ability of *Trendsetter* to correctly classify simulated sweeps of differing selection strengths, scenarios that include extensive missing data, and settings of realistic population size changes. We compare the accuracy and robustness of *Trendsetter* to other powerful methods designed to localize sweeps in single populations such as evolBoosting (Lin et al., 2011), S/HIC (Schrider and Kern, 2016b), and diploS/HIC (Kern and Schrider, 2018), and exclude complementary approaches developed to isolate sweep signals using data from multiple populations (*e.g.*, SWIF(r); Sugden et al., 2018).

#### Detecting and classifying selective sweeps

We trained *Trendsetter* with a linear (*d* = 2) trend filter penalty on data simulated under a constant-size demographic model as described in section *Materials and Methods*. We obtained optimal values for λ_1_ and λ_2_ through ten-fold cross validation. We first examined whether probability calibration was required for *Trendsetter*, and the reliability curves in Figure S2 suggest that no further calibration is needed. Based on this trained classifier, we are able to correctly classify 81.9% of hard, 97.1% of neutral, and 78.3% of soft sweep scenarios (Figure 2). Of the misclassified soft sweeps scenarios, 15.5% are misclassified as hard sweeps, and 6.2% are misclassified as neutral. We compared the performance of *Trendsetter* against several existing classification methods, where each method was modified to a three-class classification system (Lin et al., 2011; Schrider and Kern, 2016b; Kern and Schrider, 2018). Note that the native state of evolBoosting is two classes, whereas S/HIC and diploS/HIC employ five classes by default. We will examine classification ability of *Trendsetter* with five classes later in this subsection, allowing for it to be directly compared to the native states of S/HIC and diploS/HIC. From these simulated scenarios, all methods had comparable ability to detect and classify sweeps and to differentiate between hard and soft sweeps (Figures 2, 3, and S3).

**Figure 2:**
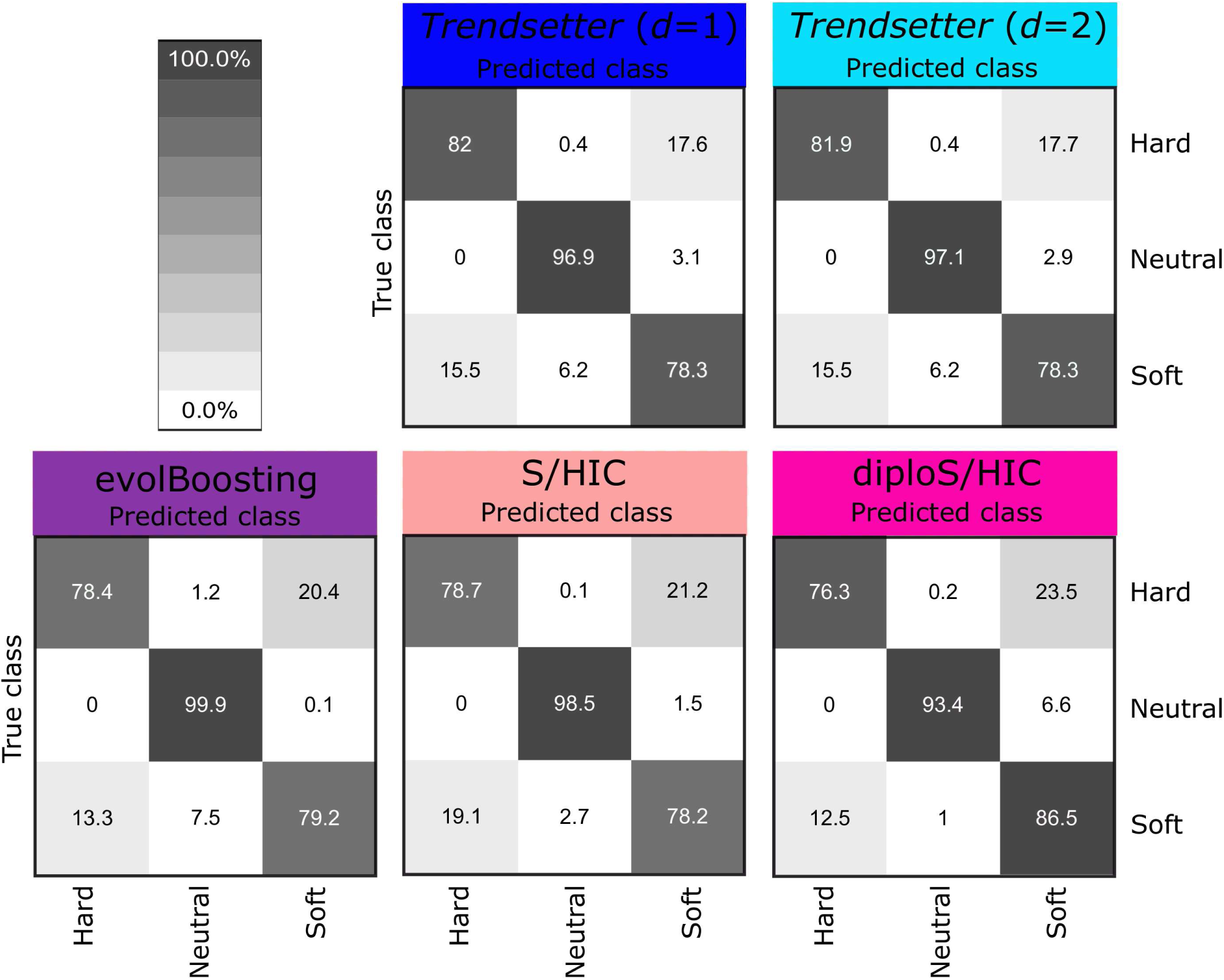
Confusion matrices comparing classification rates of *Trendsetter* with constant (*d* = 1) and linear (*d* = 2) trend penalties, evolBoosting, S/HIC, and diploS/HIC for simulations under a constant-size demographic history and selection coefficients for sweep scenarios drawn uniformly at random on a log scale of [0.005, 0.5]. All methods were trained with three classes: neutral, hard sweep, and soft sweep.

**Figure 3:**
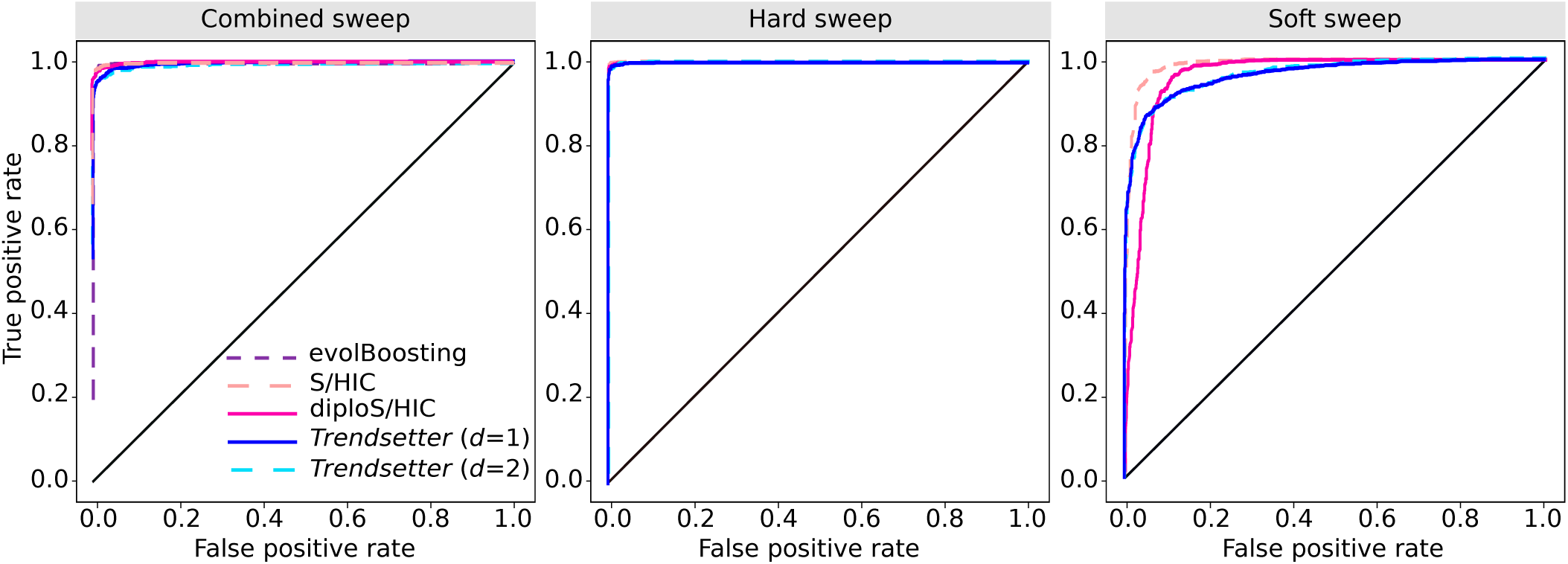
Receiver operating characteristic curves comparing the powers of various methods to distinguish sweeps from neutrality. (Left) Powers to differentiate sweeps from neutrality, by comparing the combined probability of any sweep (hard or soft) under equally-mixed hard and soft sweep simulations with the same probability under neutral simulations. (Middle) Powers to differentiate hard sweeps from neutrality, by comparing the probability of a hard sweep under hard sweep simulations with the same probability under neutral simulations. (Right) Powers to differentiate soft sweeps from neutrality, by comparing the probability of a soft sweep under soft sweep simulations with the same probability under neutral simulations. All simulations were performed under a constant-size demographic history with selection coefficients for sweep scenarios drawn uniformly at random on a log scale of [0.005, 0.5]. All methods were trained with three classes: neutral, hard sweep, and soft sweep.

By examining the values of the regression coefficients for each summary statistic, we can identify the relative importance of each statistic as well as the spatial distribution modeled. Specifically, summary statistics will tend to be more important when their regression coefficients are of larger magnitudes than other statistics. Moreover, the spatial distribution of the regression coefficients for a particular summary statistic calculated for a specific class should yield a curve, with summaries important for detecting sweeps likely exhibiting a sharp increase in magnitude near the site (central SNP) under selection (see schematic in Figure 1). These sharp peaks are the result of the combination of lasso and trend filter penalties that Trendsetter employs when fitting a regression model. If the value of a regression coefficient is reduced to zero, neighboring regression coefficients are also more likely to be zero. In contrast, the values of regression coefficients in regions of importance will be constrained by the higher values of neighboring coefficients. Figure 4 depicts the regression coefficients for *H*_12_ and the number of haplotypes *N*_haps_ under both constant (*d* = 1) and linear (*d* = 2) trend filter penalties as a function of the class and SNP position, with *Trendsetter* trained on a range of selection strengths *s* ∈ [0.005, 0.5]. We can see that number of haplotypes is clearly a less important statistic, with regression coefficients exhibiting low magnitudes at the peaks. The likely reason for this lack of importance is that, conditional on the number of SNPs, the number of distinct haplotypes will likely be narrowly constrained. In contrast, *H*_12_ played a large role in distinguishing between sweeps and neutrality, with its peak reaching the greatest magnitude, likely due to sweeps skewing the distribution of haplotype frequencies and thereby having a large influence on the *H*_12_ statistic. We also notice that some regression coefficients (such as *H*_1_ in Figure S4) tend to increase in magnitude toward the beginning or end of the analyzed region. This phenomenon may be due to the value of the coefficient only being constrained from a single direction, rather than both directions. We explore whether these upticks or downticks in regression coefficients at the ends of the analyzed region affect the accuracy of *Trendsetter* in the *Discussion* section.

**Figure 4:**
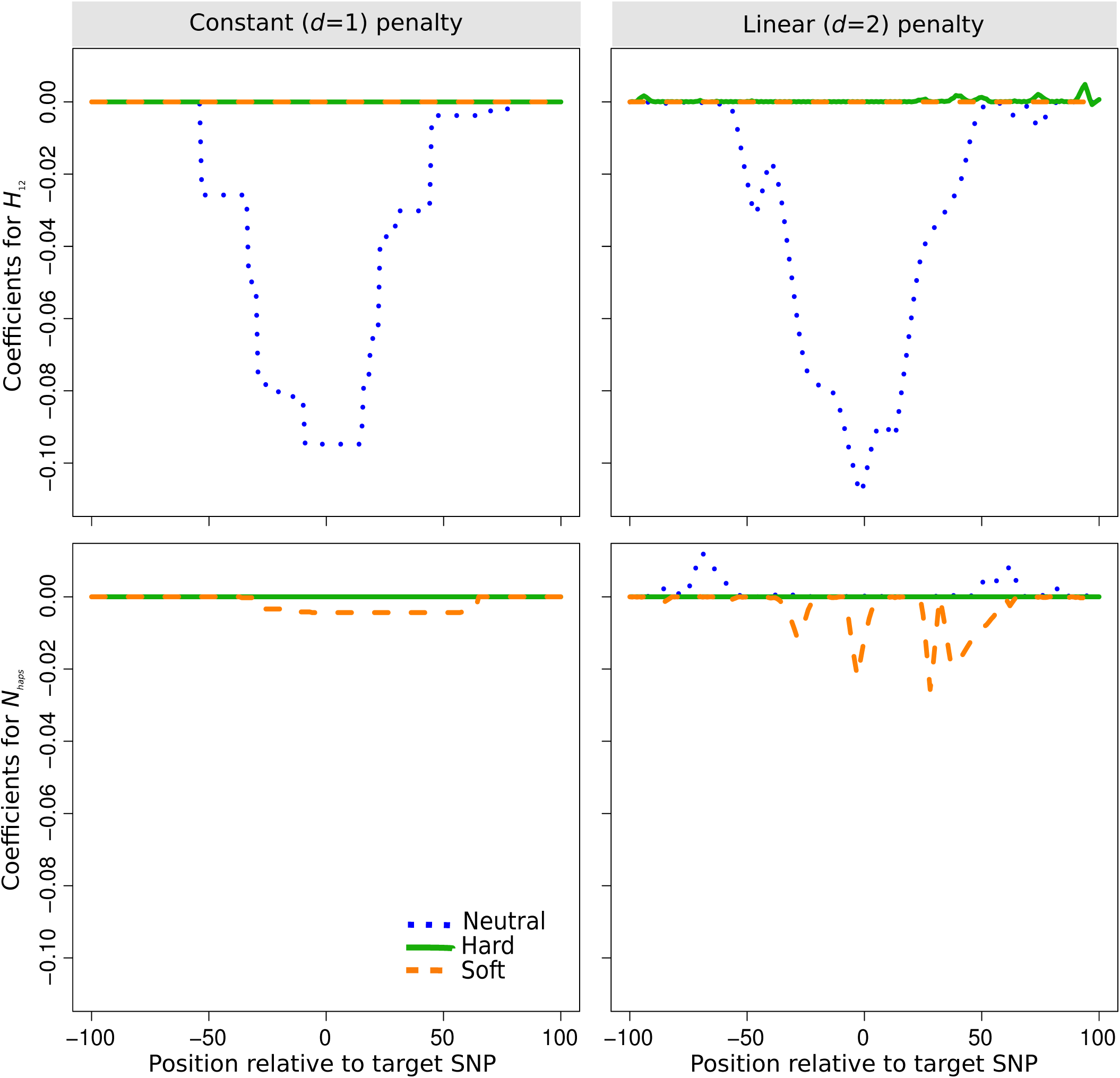
Spatial distributions of regression coefficients (*β*s) in neutral, hard sweep, and soft sweep scenarios for summary statistics *H*_12_ and number of distinct haplotypes *N*_haps_, for *Trendsetter* applied with constant (*d* = 1) and linear (*d* = 2) trend penalties. *Trendsetter* was trained on simulations with selection strength *s* ∈ [0.005, 0.5] sampled uniformly at random on a log scale. Note that the distributions of regression coefficients for both summary statistics are plotted on the same scale, thereby making the distribution of *N*_haps_ difficult to decipher as its magnitude is small relative to *H*_12_.

In a similar manner to fitting a linear (*d* = 2) trend penalty, we trained *Trendsetter* with a constant (*d* = 1) trend penalty that resulted in similar classification performance as when we trained under the linear penalty (Figures 2 and 3). This overall similarity in classification rates between constant (*d* = 1) and linear (*d* = 2) trend filtering is reflected in their similar distributions (Figure 4), with comparable relative importance levels, magnitudes, and spatial distributions of regression coefficients. Interestingly, the linear penalty is better at localizing the SNP closest to the site of selection relative to the constant penalty, based on the regression coefficients for *H*_12_ (Figure 4). Because of their similarity in performance, our discussion will be based on linear trend filtering (*d* = 2), unless otherwise specified.

We expect a disparity in the power of *Trendsetter* to detect sweeps resulting from different selection strengths *s*. The selection strength of test simulation sets strongly influences its hard sweep classification rates (Figure S5), in that simulations of strong hard sweeps are classified correctly more often than moderate hard sweeps. Further, from the curves displayed in Figure S6, we find that *Trendsetter* exhibits equal power in differentiating between neutrality and soft sweeps, regardless of the selection strength. This pattern is also reflected in Figure S5, which indicates that selection strength does not lead to substantial differences in mis-classification rates of soft sweeps for the selection strengths that we have considered—though *Trendsetter* as well as other approaches would likely have little ability to detect and classify soft sweeps from sufficiently weak selected alleles.

In contrast to the models we have considered so far, both S/HIC and diploS/HIC include two other classes in their native states, so that in addition to classes representing neutrality, hard sweeps, and soft sweeps, there are also classes representing regions that are linked (or nearby) to hard sweeps and linked to soft sweeps. The motivation for including these classes was to increase robustness of these methods to soft shoulders (Schrider et al., 2015; Schrider and Kern, 2016b; Kern and Schrider, 2018). We observe a slight increase in the mis-classification of linked-hard regions as soft sweeps (Figure S7) when we test simulations containing linked sweeps using *Trendsetter* trained to differentiate among three classes (neutrality, hard sweeps, and soft sweeps). We next chose to test whether incorporating additional (linked-hard and linked-soft) classes will increase *Trendsetter*’s robustness to soft-shoulders. Under this five-class model, the spatial distributions of regression coefficients for linked-sweep regions are modeled distinctly from sweep regions (Figure S8). Although *Trendsetter*’s ability to distinguish between hard sweeps and linked-hard regions is limited, we show that our mis-classification of linked-hard regions as soft sweeps is not dramatically different from that of S/HIC (Figures S9-S11). Because S/HIC and diploS/HIC were designed to include linked-sweep classes, we test whether including these classes alters their classification rates under confounding factors in the *Missing data* subsection of *Results*.

#### Influence of population history

Populations tend not to maintain constant sizes, with sizes instead fluctuating over time (Graciá et al., 2015; Osborne et al., 2016; Sherry, 2018). For example, it is widely accepted that global human populations have undergone different recent demographic events, such as more rapid expansions and more extreme bottlenecks in European and Asian populations when compared to Africans (Gravel et al., 2011; Tennessen et al., 2012). However, population size changes alter local genomic diversity, and can mimic signatures of selective sweeps (Galtier et al., 2000; Stajich and Hahn, 2005). It is therefore important to assess the effects of population size change on method performance.

We trained and tested *Trendsetter* on data simulated under realistic demographic models with recent population bottlenecks and expansions that are consistent with genetic variation observed in empirical human data. In particular, we generated simulation and training data from inferred human demographic parameters (Terhorst et al., 2017, see *Materials and Methods*). In general, *Trendsetter* performs well when trained and tested on realistic demographic histories (Figure 5). Simulations of African populations (LWK and YRI) showed the lowest rates of misclassification (Figure 5), likely due to their larger effective sizes and therefore greater neutral haplotypic diversity (Tenesa et al., 2007). Overall, the classification rates of simulations using *Trendsetter* with constant (*d* = 1) trend filtering (Figure S12) are virtually identical to those under linear (*d* = 2) trend filtering (Figure 5). Additionally, classification rates appear to be correlated with effective population size (Figure S13), with larger effective sizes such as in Africans leading to a greater percentage of correctly classified simulations. This trend with effective size is expected, due to the positive correlation of haplotypic diversity with effective population size.

**Figure 5:**
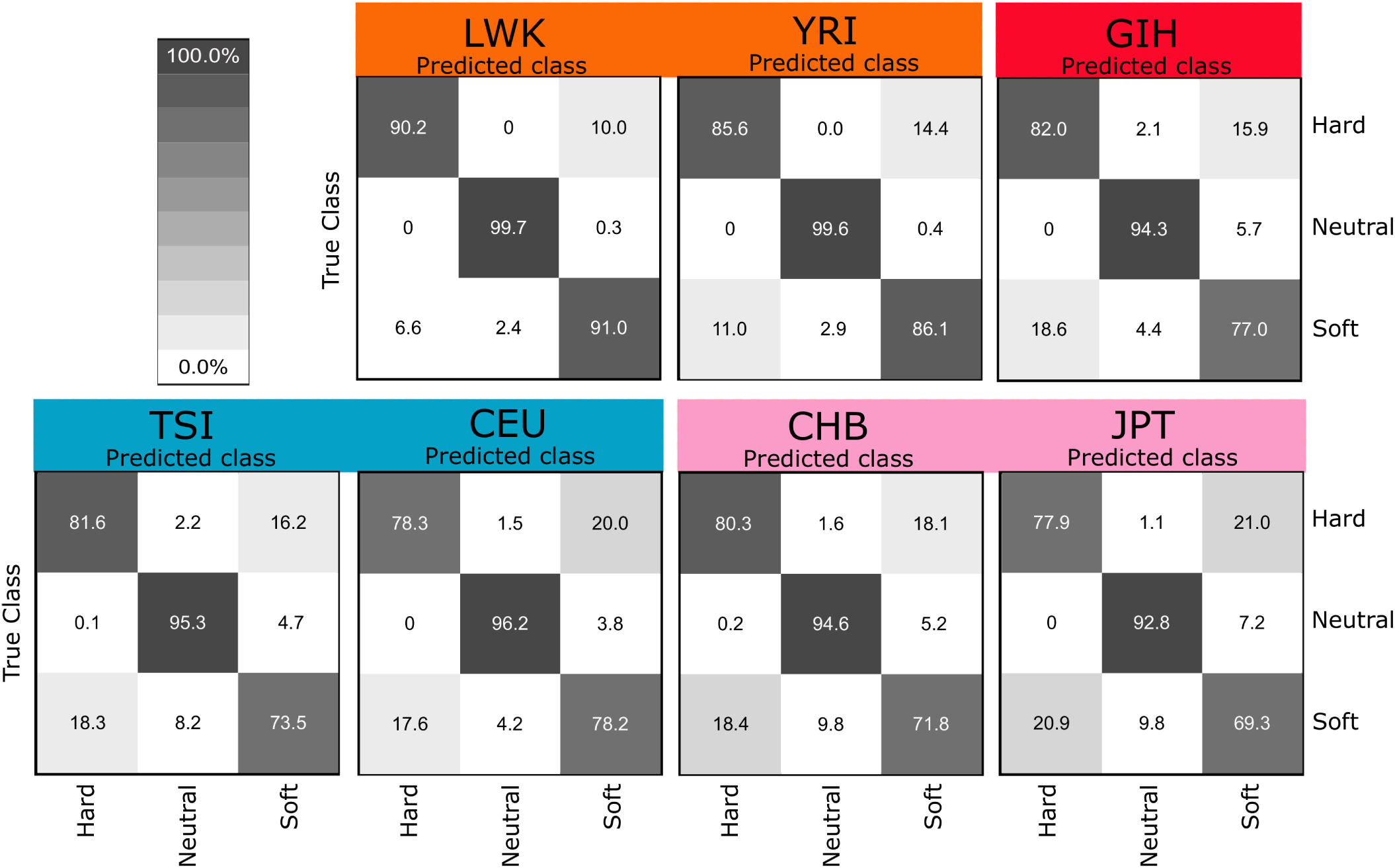
Confusion matrices comparing classification rates of *Trendsetter* with a linear (*d* = 2) trend penalty under demographic parameters estimated (Terhorst et al., 2017) from African (LWK and YRI), South Asian (GIH), European (TSI and CEU), and East Asian (CHB and JPT) populations. Selection coefficients for sweep scenarios were drawn uniformly at random on a log scale of [0.005, 0.5].

Though all classifiers performed well under diverse models of population size change, demographic misspecification (*i.e.*, testing on a population history that is different from the one that was used to train the classifier) leads to high misclassification rates if the training and test demographic histories are highly different (Figure S14). We chose to compare classification rates with demographic misspecifications for populations with similar (*e.g.*, CHB vs. GIH) and different (*e.g.*, LWK vs. CEU) histories. For all methods tested, when demographic histories of populations are similar, classification rates are not dramatically affected. However, when training on a history without a bottleneck (LWK) and testing on one with a bottleneck (CEU), *Trendsetter* (as well as evolBoosting) classified neutral regions as soft sweeps, whereas S/HIC and diploS/HIC classified soft sweeps as hard. Interestingly, by training *Trendsetter* with a combination of simulations conducted under specifications for several diverse demographic histories, we are able to improve classification rates for all test populations when demographic history is misspecified (Figure S15), and a similar performance rescue would be expected for evolBoosting, S/HIC, and diploS/HIC. As illustrated in this experiment, increasing the range of simulation parameters to reflect a more general demographic history may be desirable when training classifiers in populations for which the demographic history is not well studied.

#### Effect of sample size

Though under ideal scenarios there will be sufficient resources for studies to produce large quantities of high-quality sequence data, this is not always the case. Instead, studies may often have access only to datasets with relatively small sample sizes. The sample sizes of simulated data used to train *Trendsetter* should match that of the empirical dataset in a particular study. In our simulation examples, we evaluated the performance of *Trendetter* on a modest sample size of 50 diploid individuals. Here, we explore whether an increase or decrease in the sample size would substantially affect classification rates of *Trendsetter*, and find that the sample size does not have a great effect on classification rates. In particular, for situations in which we have half the sample size of 25 diploids (Figure S16), correct classification of hard sweep, soft sweep, and neutral scenarios was almost identical to samples of 50 diploids (Figure 2), with a slight decrease in the correct classification of hard sweeps. When we instead use a small sample of 10 diploid individuals, *Trendsetter* shows a slight decrease in correct classification rates for all classes, although it is not a dramatic difference from a sample 10 times larger (Figure S16).

Differences in sample sizes may have more of an effect on classification rates when sampled populations have gone through recent expansions or bottlenecks as experienced by human populations. For our original analyses, we sampled 50 diploid individuals from each population. To test the effect of sample size on the classification rates for a population known to have gone through a strong bottleneck (CEU) as well as no bottleneck (LWK), we trained and tested models with 100 and 25 diploid individuals for LWK and CEU demographic histories (Terhorst et al., 2017). We find that there is no appreciable difference in the classification rates between sample sizes (Figures 5 and S17).

#### Common confounding factors

Removal of low quality genomic regions is necessary when scanning empirical genomic data for selective sweeps. Depending on the stringency of filtering, this process can lead to large fractions of the genome as unclassifiable to avoid biasing scans of selection (*e.g.*, Kelley et al., 2006; Schrider and Kern, 2016b). However, it would instead be ideal if such regions could still be robustly classified despite large percentages of missing sites. We therefore chose to investigate the robustness of *Trendsetter* to excessive levels of missing segregating sites (see *Materials and Methods*). Substantial missing data in the test datasets did not significantly alter the *Trendsetter* classification rates, whereas evolBoosting, S/HIC, and diploS/HIC incorrectly classified a large percentage of simulations, including neutral simulations as hard or soft sweeps (Figures 6 and S18). Though we observed that missing data increased the mis-classification rate of soft sweeps with *Trendsetter*, these soft sweep simulations tend to be classified as neutral regions (Figure S18). Therefore, *Trendsetter* is more conservative than other comparable approaches under settings with large amounts of missing data without explicitly accounting for the distribution of missing data. The inclusion of linked-sweep classes in the model leads to most hard sweep, soft sweep, and neutral simulations with missing data to be misclassified as either linked hard or linked soft (Figures S19-S21). It should be noted that a linked-sweep classification is regarded as neutral and inclusion of linked classes leads to an increase in the performance of the method. Importantly, however, S/HIC and diploS/HIC trained with linked-sweep classes misclassify neutral simulations with missing data as soft sweeps 23.7% and 18.5% of the time, respectively (Figure S19).

**Figure 6:**
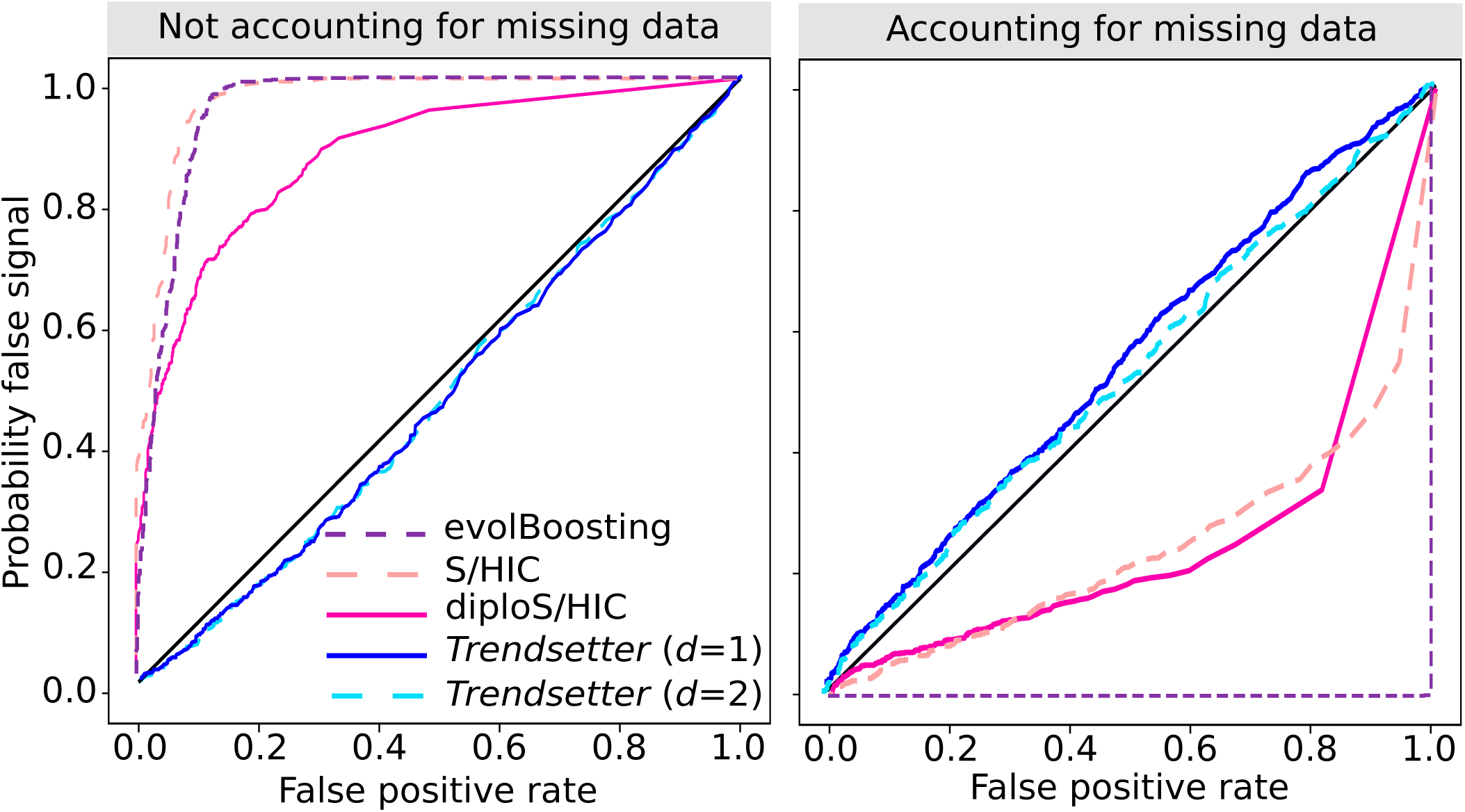
Probability of misclassifying neutral regions with extensive missing data as a sweep for various methods, under a constant-size demographic history. Each panel compares the combined probability of any sweep (hard or soft) under simulations with missing data (probability of false signal) to the same probability under neutral simulations (false positive rate) for scenarios in which missing data is (right) or is not (left) accounted for when training a classifier to compute the probability of a false signal. All methods were trained with three classes: neutral, hard sweep, and soft sweep.

The sensitivity of evolBoosting, S/HIC, and diploS/HIC is due to their reliance on summary statistics computed over large physical distances, and certain summaries, such as Tajima’s *D* or the number of distinct haplotypes, may be heavily affected by missing genomic regions. It should be noted that because we randomly removed chunks of data from simulated replicates, it is possible that data was by chance not removed from the center of simulations under neutral scenarios due to their large number of segregating sites relative to sweep settings. To address this potential issue, we randomly removed 30% of the SNPs within the central 1010 SNPs for each neutral replicate simulation and applied *Trendsetter* to the central 1010 SNPs after filtering, thereby mimicking the application of *Trendsetter* in a genomic region with extensive missing data. We find that *Trendsetter* retains its high robustness even under this scenario (Figure S22).

However, as indicated by Kern and Schrider (2018), it is also possible to train classifiers with simulations that model the distribution of missing data to account for this confounding factor. To test this idea, we trained *Trendsetter*, evolBoosting, S/HIC, and diploS/HIC on simulations with substantial missing data (*i.e.*, 30% of segregating sites missing as described in *Materials and Methods*). Accounting for the distribution of missing data when training the classifiers rescued the accuracy of all methods under this scenario, and also led to a slight boost in the overall classification rates for *Trendsetter* (Figures 6 and S23).

In addition to missing data, background selection is a ubiquitous factor (*e.g.*, McVicker et al., 2009; Comeron, 2014) that can leave similar genomic signatures as selective sweeps (Charlesworth, 2013; Nicolaisen and Desai, 2013), and which has been demonstrated to mislead sweep-detection approaches (*e.g.*, Huber et al., 2016). We examined two different scenarios of background selection—one in which a single centrally-located 11 kb protein-coding gene with strongly-deleterious alleles arising continuously is flanked by non-coding genomic regions (denoted *Central gene BGS*), and another in which potentially functional genomic regions are scattered throughout a 1.1 Mb genomic region, with the spatial distribution and distribution of fitness effects of deleterious alleles inspired by their respective distributions in humans (denoted *Empirical-based BGS*) as described in *Materials and Methods*.

*Trendsetter*, S/HIC, and diploS/HIC are relatively robust to both forms of background selection (Figures 7 and S24), with *Trendsetter* demonstrating slightly better performance than S/HIC and diploS/HIC as it almost always classifies background selection as neutral. In contrast, S/HIC and diploS/HIC sometimes classify regions of background selection as soft sweeps, and evolBoosting almost always classifies background selection as a soft sweep. It is important to note that S/HIC and diploS/HIC likely classify some background selection simulations as soft sweeps because we have not included the two linked-sweep classes that they generally employ. The inclusion of such classes would probably lead to such regions being classified as a linked sweep, which should be regarded as neutral. Moreover, the poor performance of evolBoosting is due to Tajima’s *D* being the feature of greatest importance for all windows in the trained classifier, which can be mislead by background selection as it can generate distortions in the site frequency spectrum that are similar to sweeps (Charlesworth, 2013; Nicolaisen and Desai, 2013).

**Figure 7:**
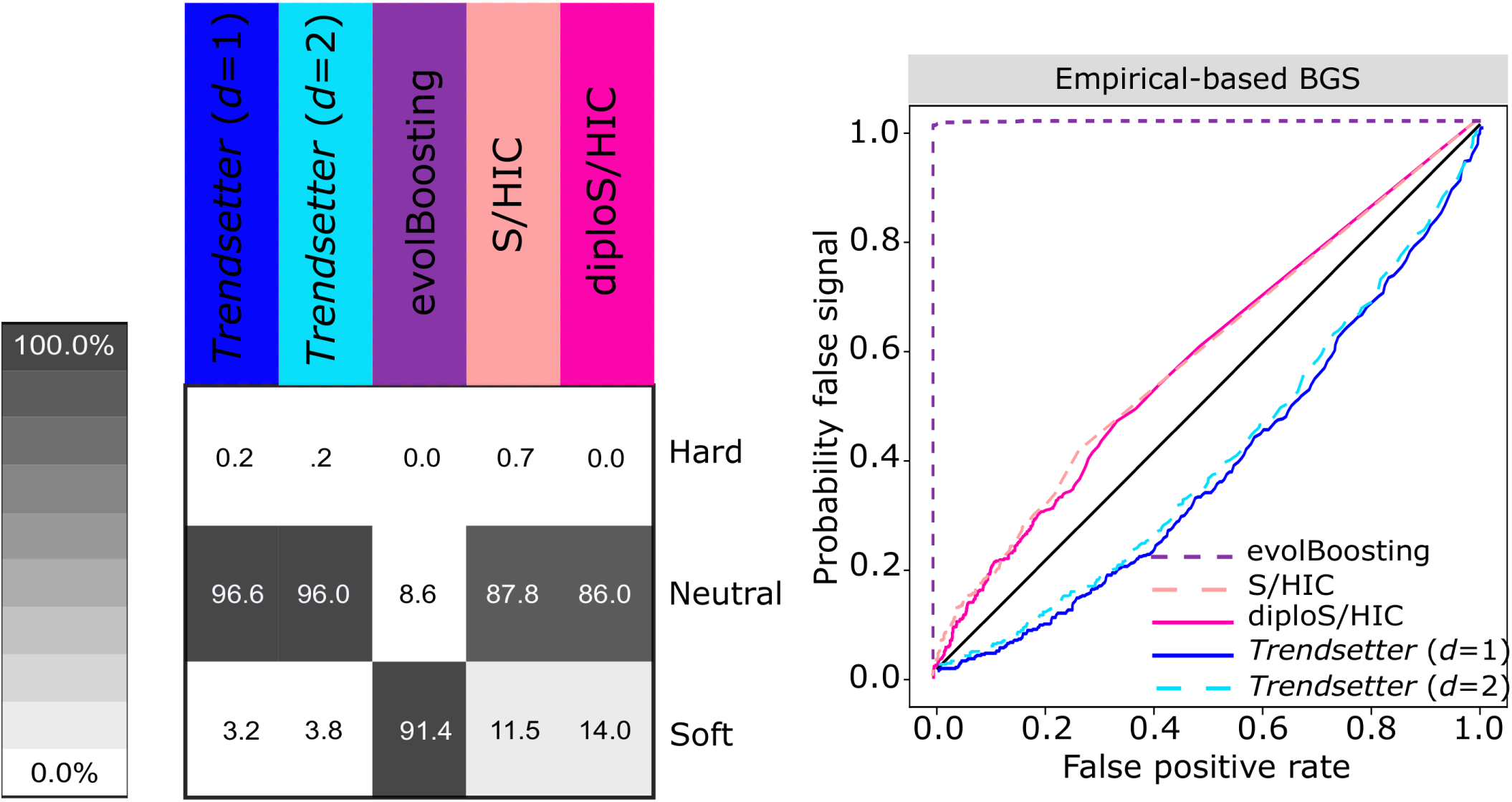
Robustness of mis-classifying genomic regions undergoing background selection for various methods, under a constant-size demographic history and background selection parameters (number, lengths, and distribution of functional sites as well as distribution of fitness effects) drawn from a distribution based on human data (see *Materials and methods*). (Left) Classification rates for regions evolving under background selection. (Right) Probability of misclassifying regions evolving under background selection, by comparing the combined probability of any sweep (hard or soft) under simulations with background selection (probability of false signal) to the same probability under neutral simulations (false positive rate). All methods were trained with three classes: neutral, hard sweep, and soft sweep, with selection coefficients for sweep scenarios drawn uniformly at random on a log scale of [0.005, 0.5].

We also examined a modification to the *Central gene BGS* scenario, by decreasing the recombination rate in the centrally-located 11 kb genic region by 100 fold (denoted *Central gene BGS with low recombination*), which is meant to simulate a massive reduction in haplotypic diversity within the central genic region as strongly-deleterious mutations arise in the region. We find that classification rates for all methods decrease by only a few percentage points under this scenario (Figure S24). This set of background selection simulations demonstrates that *Trendsetter*, S/HIC, and diploS/HIC are robust to typical as well as strong background selection, and robustness of all methods could likely be improved by including an additional class for background selection (Schrider and Kern, 2016b).

Another potential confounding factor is the fluctuation of recombination rate across the genome, as recombination rate changes can influence haplotypic diversity and therefore impact sweep detection. To examine the influence of this factor on sweep detection and classification, we simulated genomic regions with lower (*r* = 5 × 10^-9^) and higher (*r* = 5 × 10 ^-8^) recombination rates compared to the recombination rate (*r* = 10^-8^) used to simulate the training data. We find that both increasing and decreasing recombination rates leads to high misclassification rates (Figure S25). In general, regions of lower recombination typically lead to an increase in the rate of misclassifying soft sweeps as hard sweeps across all compared methods. In contrast, higher recombination rate regions lead to an increase in rates of mis-classifying hard sweeps as soft sweeps across all methods. Moreover, *Trendsetter* and evolBoosting also have an elevated rate of misclassifying soft sweeps as neutral regions. These results suggest that accounting for recombination rate variation when training a classifier is highly important, as not considering a range of recombination rates could lead to mis-classification of the types of identified sweeps.

#### Application to empirical data

Global human populations have encountered a number of diverse environments in their past, likely leading to various adaptive pressures experienced across populations (Sabeti et al., 2006; Hancock et al., 2008). For this reason, we sought to identify genomic regions that are likely candidates for recent selective sweeps in different populations. Because our results on recombination rate changes on simulated data indicated that *Trendsetter* is not robust to recombination rate variation when it is not directly accounted for in the training step, we simulated training replicates in which recombination rates were drawn from an exponential distribution with mean 10^-8^and truncated at three times the mean as in Schrider and Kern (2017). We also show that classifiers are reasonably-well calibrated for demographic histories of all populations that we consider in our empirical analysis after training with recombination rate variation (Figure S26).

Classification of populations from the 1000 Genomes Project (The 1000 Genomes Project Consortium, 2015) showed in general that recent hard sweeps are relatively rare, as has been previously demonstrated in humans and other species (*e.g.*, Garud et al., 2015; Schrider and Kern, 2017). For our empirical scan we classify every fifth autosomal SNP, beginning from the center SNP in the 101st window (505th SNP) as described in the *Application to emprical data* subsection of *Materials and Methods*. We compute the fraction of autosomes classified as a certain class as the fraction of classified SNPs belonging to that class. Between 0.00 and 1.83% of each chromosome was classified as a hard sweep, and between 4.61 and 12.95% was classified as soft when we trained *Trendsetter* using demographic parameters inferred by Terhorst et al. (2017) (Tables S1-S3). *Trendsetter* also detected genes previously identified as hard sweeps, such as *EDAR* in CHB (Bryk et al., 2008). Figure S27 shows the probability of a hard sweep under *Trendsetter* across the region on chromosome 2 surrounding *EDAR* in the seven global populations considered, and displays a clear peak under the selected gene *EDAR* in the East Asian (CHB and JPT) populations. By examining the values of the summary statistics calculated in the region containing *EDAR* for the Han Chinese (CHB) population (Figure S27), we see that there are clear decreases in the values of 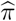, number of haplotypes *N*_haps_, and *H*_2_*/H*_1_, as well as increases in the values of *H*_1_ and *H*_12_, providing support for the strong hard-sweep classification in this region (Figure S27). We also find that the *LCT* gene, which is classified as a soft sweep in the CEU population, shows similar patterns of summary statistics in the region of selection (Figure S28). Although the region surrounding *LCT* has been previously classified as a hard sweep (Peter et al., 2012), a more recent study by Schrider and Kern (2016b) also classifies the region surrounding *LCT* as soft. Moreover, we identify as soft sweeps many genes previously hypothesized to be under positive selection, such as *TRPV6* (Figure S29), *PPARG*, and *EPHB6* (Akey et al., 2004). *TRPV6* was also discovered by Peter et al. (2012), but was not classified as either hard or soft.

We also uncover a number of novel candidate sweeps. For many genes classified as positively selected in a population, these genes are also classified as under positive selection in other human populations. Among these are cancer-related genes, such as *BRCA1* and *FBXW7*. *BRCA1* was classified as a soft sweep in East Asian (CHB and JPT), South Asian (GIH), and European (CEU and TSI) populations (Figure S30). The distribution of summary statistic values used to classify this region also display expected sweep patterns (Figure S30). Moreover, *FBXW7*, a tumor suppressor gene in which mutations are associated with colorectal, ovarian, and liver cancers (Jardim et al., 2014), was classified as a soft sweep in six (LWK, GIH, TSI, CEU, CHB, and JPT) out of the seven populations that we evaluated (Figure S31). Interestingly, Schrider and Kern (2017) also reported that a large number of genes they determined to be influenced by a sweep have been associated with cancer. Furthermore, there exists prior evidence of positive selection acting on cancer-related genes, such as *BRCA1* (Lou et al., 2014), which may help explain the high percentage of cancer-related genes flagged as candidate sweep targets by *Trendsetter*.

In addition to signals over specific genes, we observe in general that regions classified as a sweep tend to be shared across populations. Specifically, we find that genomic regions classified as either hard or soft sweeps tend to be classified as the same sweep class in other populations. To quantify this observation, we measure the extent to which sweeps signals in one population are also found in other populations. In particular, we computed the fraction of non-overlapping 10 kb genomic segments classified as a soft (hard) sweep in a given population that are also classified as a soft (hard) sweep in another population. We find that populations share more soft sweeps with populations from the same geographic region than with populations from other regions (Figure 8), most likely resulting from shared ancestry rather than convergent evolution. The African populations (LWK and YRI) form a cluster of shared sweeps as do the East Asian (CHB and JPT) and separately, European (TSI and CEU) populations. European populations also form a sharing cluster with the South Asian population GIH. Although the proportions are much higher when quantifying shared hard sweeps (Figure S32), the patterns of sweep sharing are similar to that of soft sweeps (Figure 8) and mimic the sharing of haplotypes across globally-distributed human populations observed by Conrad et al. (2006).

**Figure 8:**
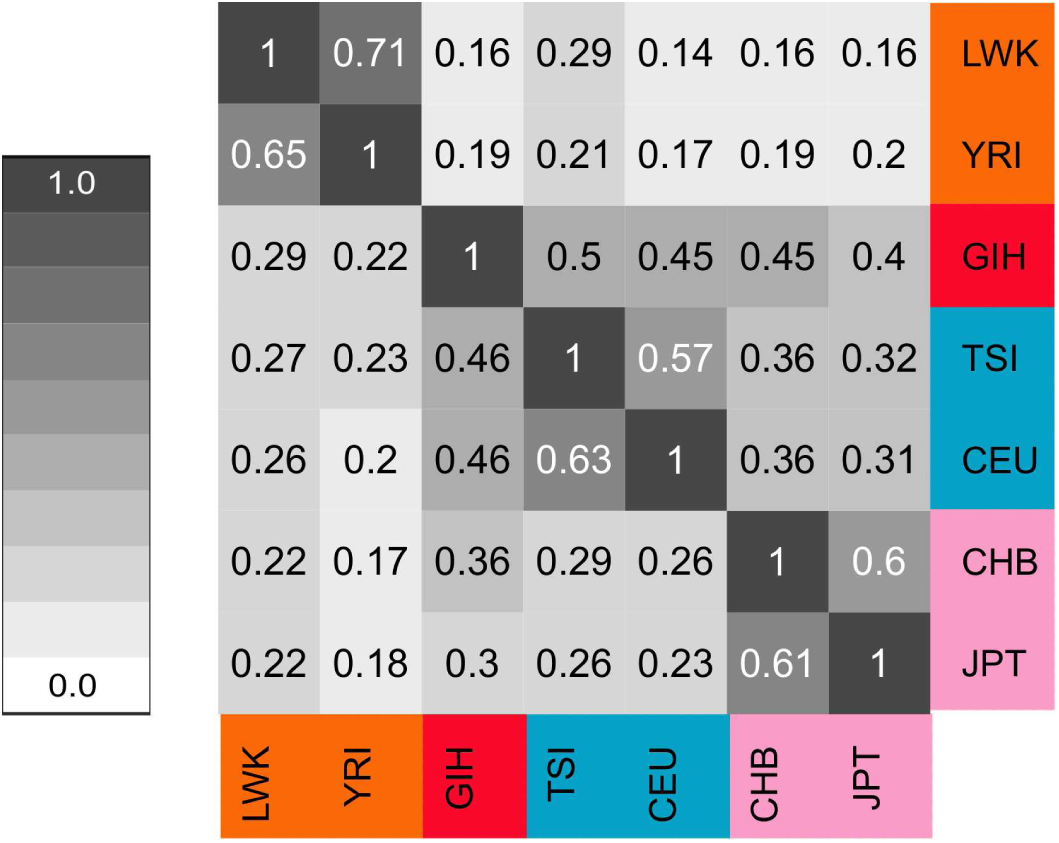
Heatmap representing the sharing of soft sweep classifications across worldwide human populations. The cell at row *j* and column *k* represents the proportion of non-overlapping 10 kb genomic segments classified as a soft sweep in the population at row *j* that are also classified as a soft sweep in the population at column *k*. By definition, this heatmap is asymmetric.

### Discussion

In this article we demonstrated the ability of *Trendsetter* to localize and classify selective sweeps from the spatial distribution of summary statistics in the genome. In its current form, *Trendsetter* uses information from six different summary statistics to differentiate among three classes—neutrality, hard sweeps, and soft sweeps. Based on this formulation of *Trendsetter*, we found that it is resistant to common issues such as missing genomic segments (Figures 6 and S18) and background selection (Figures 7 and S24). This robustness to such confounding factors is likely due to its reliance on haplotype-based statistics such as *H*_1_, *H*_12_, and *H*_2_*/H*_1_ (Garud et al., 2015), to its use of SNP-based windows for calculating summary statistics, and to the use of the spatial distribution of each summary statistic. Other approaches that rely on statistics that emphasize the number of segregating sites or the site frequency spectrum, such as Watterson’s 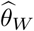 or Tajima’s *D*, in a window may have higher power to detect sweeps, but also exhibit higher misclassification error rates, leading to regions harboring extensive missing data or undergoing background selection to be mistaken as candidate sweep regions (Figures 6, 7, and S18). However, as we demonstrated in Figure S23, this lack of robustness to, for example, missing data may be remedied by training models with simulations including missing data.

Flexibility in the choice of summary statistics allows *Trendsetter* as well as other complementary approaches (Lin et al., 2011; Schrider and Kern, 2016b; Kern and Schrider, 2018) to be easily applied to a number of settings, and for this reason, particular choices of summary statistics for other approaches may also lead to greater robustness to confounding factors but with likely power trade-offs. *Trendsetter*’s ability to correctly classify sweeps and distinguish sweeps from neutrality increases when we trained a model with S/HIC and diploS/HIC-specific statistics calculated in 110 contiguous windows each of length 10 kb (Figure S33). We chose this large number of windows so that we could learn the spatial distribution of each summary statistic. However, if we normalize each statistic across the set of windows it is calculated (as in S/HIC and diploS/HIC; Schrider and Kern, 2016b; Kern and Schrider, 2018), mis-classification between hard and soft sweeps increases (Figure S33, right column).

The types of summary statistics employed by *Trendsetter* contribute to the reason for its robustness to missing data. We tested whether training *Trendsetter* with the complementary sets of summary statistics as used by S/HIC or diploS/HIC would affect *Trendsetter*’s classification rates under missing data. In contrast to the patterns displayed by S/HIC (Figure S18), we observed a larger percentage of mis-classifications toward soft sweeps rather than toward hard sweeps, when we use non-normalized versions of S/HIC-specific statistics (Figure S34). If we chose to instead normalize statistics, then mis-classification to hard sweep increases (Figure S34). Moreover, these latter results mirror those observed for S/HIC (Figure S18), which uses the identical normalization procedure for summary statistics computed across a genomic region. Similarly, we observe that simulations with missing data tend to be mis-classified as hard when *Trendsetter* employs normalized versions of diploS/HIC statistics (Figure S34), computed in an analogous manner with 110 contiguous windows each of length 10 kb.

As in Figure S23, using the set of S/HIC and diploS/HIC statistics combined with training under missing data would likely lead to a powerful classifier that is also robust to missing data. Interestingly, when we also trained and tested *Trendsetter* using diploS/HIC-specific summary statistics with demographic mis-specifications as described in *Influence of population history*, we found that when *Trendsetter* is trained and tested with normalized diploS/HIC-specific statistics (Figure S35) we recapitulate the classification patterns of diploS/HIC trained and tested under demographic mis specification (Figure S14). Therefore, the choice of the set of summary statistics may have a large influence on the behavior of a sweep classifier, regardless of the diverse set of approaches (*e.g.*, random forests, neural networks, or regularized regression) employed to model the data.

We also tested the classification rates of *Trendsetter* when operating on S/HICand diploS/HIC-specific statistics for *K* = 5 classes, representing neutral, hard sweep, soft sweep, linked to hard sweep, and linked to soft sweep scenarios, used by those methods, calculated in 110 contiguous 10 kb-long windows. *Trendsetter* using *Trendsetter* -specific statistics (Figure S9-S11) exhibited comparable performance to *Trendsetter* using diploS/HICand S/HIC-specific statistics (Figure S36). Testing the classification of simulations with missing data we find that most neutral simulations missing data are classified as linked to a soft sweep (Figure S37), though there is also a large misclassification rate toward soft sweeps. We also examined the magnitude and spatial distribution of regression coefficients for *Trendsetter* using diploS/HIC-specific statistics to evaluate feature importance (Figures S38). Based on the magnitudes of the regression coefficients, we find that Fay and Wu’s *H* and Watterson’s 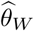 are the most informative, whereas Tajima’s *D* is among the least informative. Moreover, the peaks of the curves modeling each summary statistic tend to be narrow, and identify the location of the polymorphism closest to the selected site from the sweep classes. Importantly, we computed these summary statistics across data encompassing entire 1.1 Mb genomic regions when compared to the SNP-based summary statistics that we employed earlier where we used information across only 1010 SNPs. The SNP-based method of calculating statistics often used information from less than one-third of the 1.1 Mb genomic region.

Unphased multilocus genotype data are more widely available than phased haplotype data, as it can be difficult to phase genotypes for a number of study systems (Browning and Browning, 2011). Although it is possible and common to infer haplotypes from genotype data, this process is not error free (Browning and Browning, 2011), and these errors may have deleterious effects on downstream efforts to localize selective sweeps. However, it should still be possible to uncover and classify sweep regions without phased haplotypes (*e.g.*, Harris et al., 2018; Kern and Schrider, 2018). Substituting haplotype-based summary statistics with their unphased multilocus genotype analogues (see *Materials and Methods*), we find that *Trendsetter* can still differentiate well among hard sweeps, soft sweeps, and neutrality (Figure S39, six summary statistics). By examining the spatial distributions of regression coefficients for each summary statistic (Figure S40), we find that the inferred model relied heavily on the number of multilocus genotypes to make predictions, with the other summary statistics providing marginal information conditional on the number of multilocus genotypes. For settings in which phased haplotypes cannot be obtained, a hybrid approach of incorporating some additional summary statistics computed by diploS/HIC (*e.g.*, measures of the distribution, such as variance, skewness, and kurtosis of differences between pairs of individuals) in SNP-based rather than physical distance-based windows may aid classification. Incorporating these statistics slightly increases the overall accuracy (Figure S39; nine summary statistics) and shows similar feature importance patterns as when *Trendsetter* is trained without these statistics (Figure S41).

In some scenarios (*e.g.*, for *H*_1_ in Figure S4), the distribution of regression coefficients for particular summary statistics exhibited sudden increases or decreases in magnitude near edges of their genomic range. This phenomena may be due to the fact that the coefficients at the ends of this range are only constrained from one side by *Trendsetter*, whereas coefficients near the center are constrained on both sides. To verify that these changes in magnitude at the edges do not affect the classification rates of *Trendsetter*, we discarded five coefficients (and associated summary statistic values) at each end to make predictions after the model was trained, thereby removing these potential artifacts. We tested the model whose coefficients are depicted in Figure S4 without the first five and last five predictors for all summary statistics, and find that there is virtually no difference in classification rates from when they are included in the model (compare Figures 2 and S42).

Our experiments show no extensive difference in classification rates when we apply a constant (*d* = 1) versus linear (*d* = 2) trend filter penalty for differentiating among hard sweeps, soft sweeps, and neutrality (Figures S38, S43, S44, and S45). However, it is possible that for differentiating between other selection settings, such as in scenarios of adaptive introgression (Racimo et al., 2017) or in distinguishing between partial sweeps and recent balancing selection, the application of a linear rather than constant trend filter penalty will create a meaningful difference between classification rates. Regardless of the form of the trend filter penalty, we have shown that *Trendsetter* is flexible and has comparably high power to a number of previously-published statistical learning approaches for single populations. Moreover, the model learned by *Trendsetter* is a set of curves modeling summaries of genetic variation, and it is therefore easy to visualize the broad spatial distribution of summary statistic importance by construction.

Implementing *Trendsetter* as we have generally considered in this article with the six summary statistics *r*^2^, 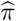, *N*_haps_, *H*_1_, *H*_12_, and *H*_2_*/H*_1_ calculated in 201 overlapping windows is unable to identify the beneficial polymorphism. Scans across simulations of hard sweeps under a constant-size demographic history with the beneficial mutation arising in the center of a 1.1 Mb region pinpoint the mean physical distance of the classified polymorphism with highest sweep probability to be 2093 bases away from the center, with the largest physical distance of 331 kb. The ability to localize adaptive regions in empirical data would also likely be affected by additional factors, such as non-equilibrium demographic history, recombination rate fluctuation, and missing data. However, incorporation of summary statistics such as iHS (Voight et al., 2006) and *nS*_*L*_ (Ferrer-Admetlla et al., 2014) may more precisely localize the SNP under selection. These statistics, although haplotype based, provide a value at every SNP and are by construction not window based, in contrast to the haplotype-based statistics employed in this article. Moreover, incorporating information from an additional population (*e.g.*, Sugden et al., 2018) would also allow us to apply powerful cross-population haplotype-based statistics such as XP-EHH (Sabeti et al., 2007) that are also based on population differentiation, and that compute a single value at each SNP. We note that large numbers of summary statistics may be provided to the model with our incorporation of a lasso penalty to help alleviate issues with over-fitting (Tibshirani, 1996). Finally, we show how *Trendsetter* can easily use any summary statistics specified by the user, which allows *Trendsetter* to be adaptable to a variety of selection scenarios users may be interested in. A Python script implementing *Trendsetter* as well as probabilities of neutral, hard sweep, and soft sweep classes for polymorphisms classified in our empirical scans can be downloaded at http://www.personal.psu.edu/mxd60/trendsetter.html.

## Acknowledgments

We thank Jonathan Terhorst for providing the SMC++ demographic history estimates from Terhorst et al. (2017), Daniel Schrider and Andrew Kern for their help with discoal, and Alexandre Harris for his assistance in preparing scripts for demographic simulations. We also thank Andrew Kern and two anonymous reviewers for their constructive feedback that helped strengthen this manuscript. This research was funded by National Institutes of Health grant R35GM128590, the Alfred P. Sloan Foundation, Pennsylvania State University startup funds, a NIGMS funded training grant on Computation, Bioinformatics, and Statistics (Predoctoral Training Program T32GM102057), and the NASA Pennsylvania Space Grant Graduate Fellowship. Portions of this research were conducted with Advanced CyberInfrastructure computational resources provided by the Institute for CyberScience at Pennsylvania State University.

